# *Vibrio cholerae* adhesin-derived peptide mediates strong pull-off forces in aqueous high-ionic-strength environments

**DOI:** 10.1101/2025.08.25.672170

**Authors:** Syeda Tajin Ahmed, Sixin Zhai, Xin Huang, Sarvagya Sulaja, Adekunle Adewole, Alisa Ioffe, Andrea D. Merg, Jing Yan, Roberto C. Andresen Eguiluz

## Abstract

In this letter, the pull-off forces of adsorbed films of four Bap1-inspired peptides in various solvents were investigated on negatively charged mica substrates using the surface forces apparatus (SFA), complemented with dynamic light scattering (DLS) for characterizing the aggregation behavior of peptides in solution. Bap1-inspired peptides consisted of the 57 amino acid wild-type sequence (WT); a scrambled version of the WT used to investigate the impact of the primary amino acid sequence in pull-off forces (Scr); a ten amino acid sequence rich in hydrophobic content (CP) of the WT sequence, and an eight amino acid sequence (Sh1) that corresponds to the pseudo-repeating sequence in the 57 AA. SFA results showed remarkable pull-off forces for CP, particularly in the presence of salts: measured pull-off forces were 26.0 ± 7.0 mN/m for no dwell-time and up to 42.0 ± 8.8 mN/m when surfaces were left in contact for 30 minutes. DLS observations indicate that salts favor large peptide aggregation for all constructs (*H*_z_ > 1 µm), as compared to milliQ (*H*_z_ ≈ 100-500 nm) water and DMSO (*H*_z_ ≈ 100 nm), resulting in heterogeneous peptide film thicknesses. This letter concludes with a comparison to the pull-off forces of mussel foot protein-inspired peptides reported in the literature.

---

Adhesion in the presence of water presents significant challenges due to water’s interference with adhesive-substrate interactions. These challenges include weakened electrostatic interactions due to ion screening,^1^ competition with water molecules and hydration layers,^2^ reduced van der Waals interactions,^3^ or oxidation.^4^ Understanding how to overcome these challenges is crucial for developing wet and salt-tolerant materials in wet environments, such as biomedical glues, marine infrastructure sealants, and a molecular understanding of biofilm formation for developing peptide-based antibacterial agents. Organisms, such as bacteria, have developed strategies to overcome challenges in the presence of water. For example, bacterial colonies rapidly cover any available surface via specific and nonspecific interactions in benthic environments or the human body. To do so, bacteria express multiple adhesive biomolecules, such as siderophores^5^ and adhesins^6^ to interact with its environment and colonize surfaces efficiently. Recently, a 57-amino-acid (AA) sequence in the Bap1 adhesin in the biofilm of *Vibrio cholerae* (Figure 1a), the causal agent of the pandemic cholera, has been identified as a critical adhesion mediator to multiple abiotic and biotic surfaces in *V. cholerae* biofilm.^7,8^ This domain has the potential to be an essential “blueprint” provided by nature for identifying new adhesive motifs based solely on canonical amino acids.

**Figure 1.**
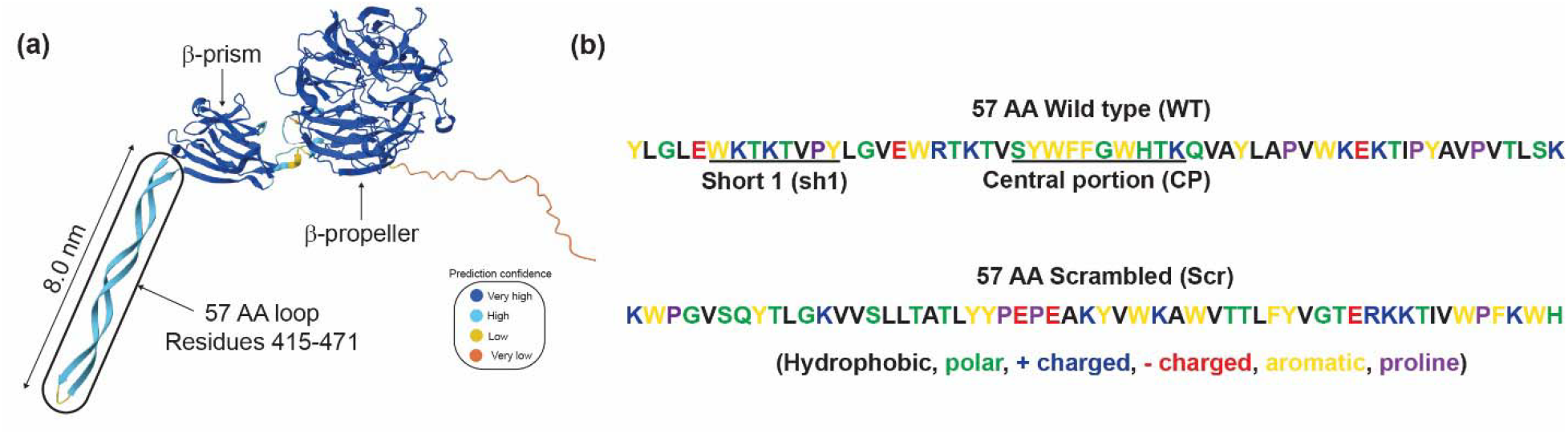
(a) 3-D representation of the *V. cholerae* adhesin Bap1, showing the main β-propeller domain, the β-prism, and the 57 AA loop relevant to this study. The β-propeller and β-prism domains have been crystalized,^7^ where the 57 AA loop structure comes from prediction of AlphaFold.^9,10^ (b) The color-coded 57 AA sequence of the wildtype (WT) loop shown in (a), and two additional peptides investigated in this study, short 1 (Sh1) and central portion (CP), as well as a scrambled (Scr) analog of the WT 57 AA.

Multiple organisms have inspired research efforts to develop wet adhesion technologies. Snails, for example, secrete various types of mucus containing tailored compositions of glycosaminoglycans, gel-forming mucins, glycoproteins, and lectins that impart lubricating properties to facilitate glide or adhesion.^11,12^ Furthermore, these mucus have shown promising results for wound healing applications.^11,13^ Studies of tick saliva or cement, which contain glycine-rich proteins chaperoned by cationic and aromatic amino acids, are yet another example of efficient and robust wet adhesin.^14^ The proposed mechanisms behind these cements’ adhesive properties are phase separation and condensate formation driven by π-π and cation-π interactions.^15^ Indeed, π-π and π-cation interactions and complementary intra- and intermolecular forces, such as hydrogen bonding, are essential, enabling strong wet cohesion and adhesion^16^ in several marine organisms.^14,17–19^ Many of these findings have originated from marine mollusks, such as mussels and barnacles. These are the organisms that have unleashed the most significant wave of wet adhesion technology promises.^20–28^ That is in part due to the abundance of catechol present in the adhesive proteins in the form of dihydroxyphenylalanine (DOPA, a posttranslational modification of tyrosine). However, given the reactivity of the catechol moiety of DOPA,^29,30^ making them more susceptible to degradation to environmental factors, other catechol-inspired, more stable moieties have been explored as potential adhesion mediators.^31,32^

In this letter, the pull-off forces, as key measurements of surface interactions, of physically adsorbed films of four Bap1-inspired peptides in various solvents were investigated on negatively charged mica substrates using the surface forces apparatus (SFA), complemented with dynamic light scattering (DLS) for characterizing the aggregation behavior of peptides in solution. Bap1-inspired peptides consisted of the 57 amino acid wild-type sequence (WT); a scrambled version of the WT (abbreviated as Scr) used to investigate the impact of the primary amino acid sequence in pull-off forces; a shorter ten amino acid sequence rich in hydrophobic content corresponding to the central portion (CP) of the WT sequence that was recently identified to be critical for lipid binding;^33^ and an eight amino acid sequence named short-1 (Sh1) that corresponds to the pseudo-repeating sequence in the 57 AA (Figure 1b, 1c, and SI S1 Bap1-inspired peptide details), all synthesized from at least two independent sources (SI S2 Bap1-peptide synthesis and characterization). Peptides were dissolved in dimethyl sulfoxide (DMSO) at a stock concentration of 150 µM and used to prepare working solutions of 1.5 µM of peptides in either aqueous solvents with high ionic strength (M9 buffer at pH 7, with a total ionic strength of 190 mM, which is a common bacteria growth medium) or in deionized water (milliQ water at pH 7, no salts), or further diluted with an organic solvent (pure DMSO) to 1.5 µM. SFA results showed remarkable bridging forces (pull-off forces) for CP, particularly in the presence of ions (M9): measured pull-off forces were 26.0 ± 7.0 mN/m (5.3 ± 1.4 mJ/m^2^) for no dwell time and up to 42.0 ± 8.8 mN/m (9.0 ± 1.7 mJ/m^2^) when surfaces were left in contact for 30 minutes. DLS observations indicate that ions present in M9 favor large peptide aggregation for all constructs (with hydrodynamic diameter, *D*_H_ > 1 µm), as compared to milliQ (*D*_H_ ≈ 100-500 nm) water and DMSO (*D*_H_ ≈ 5 nm), resulting in heterogeneous peptide film thicknesses, as measured by SFA. This letter concludes with a comparison to the pull-off forces of mussel foot protein (mfps) inspired peptides reported in the literature.

First, to quantify the pull-off forces of WT, Scr, CP, and Sh1 peptide films deposited onto mica substrates from an M9 aqueous buffer (Figure 2a), the SFA was used. Details on the SFA and the symmetric peptide depositions procedures can be found in the SI sections *S4 Surface Forces Apparatus and substrate preparation* and *S5 Peptide film deposition*. Representative SFA force-distance curves in M9 buffer, measured for WT, CP, and Sh1 are shown (Figure 2b). Scr formed large aggregates, resulting in double contacts, and it was not possible to extract pull-off forces from the force spectroscopy measurements performed with SFA, and therefore not shown in Figure 2a. All force-distance curves for WT and CP exhibited purely repulsive forces during approach, followed by an abrupt jump out of contact during retraction of the surfaces, which was distinct in magnitude from that observed for the M9 buffer alone (Figure S2). Sh1 did adsorb to the mica, forming a peptide film, but only repulsive forces were detected during approach, and no jump-out of contacts occurred. Pull-off forces (force normalized by radius of curvature required to separate the two participating surfaces), -*F/R*, for WT and CP peptides were measured to be 18.0 ± 1.0 mN/m and 26 ± 6.9, respectively, with no dwell time (total contact time of approximately 2 minutes, time it takes to complete the loading/unloading cycle). Leaving the surfaces under the maximum applied load (□ 100 mN/m) for an additional 10 min or 30 min had a marginal effect on WT films, increasing the pull-off forces to 21 ± 2 mN/m (Figure 2c). For CP, however, pull-off forces increased to 29 ± 9 mN/m and 42 ± 9 mN/m for 10 and 30 min, respectively, corresponding to an increase of 10% and 60% in the magnitude of pull-off forces. These observations suggest that most of the intra- and intermolecular interactions responsible for the adhesive and cohesive properties of the WT nanofilm were formed during the shorter contact times and remained stable. For CP, however, intra- and intermolecular interactions responsible for the adhesive and cohesive properties of the CP nanofilm appear to increase, indicative of a higher degree of conformational flexibility of the nanofilm, and providing a higher density of effective “sticky” moieties per peptide molecule. This is further evidenced by the different film thicknesses formed between WT, Sh1, and those formed by CP, with CP forming significantly larger nanofilms (Figure 2d).

**Figure 2.**
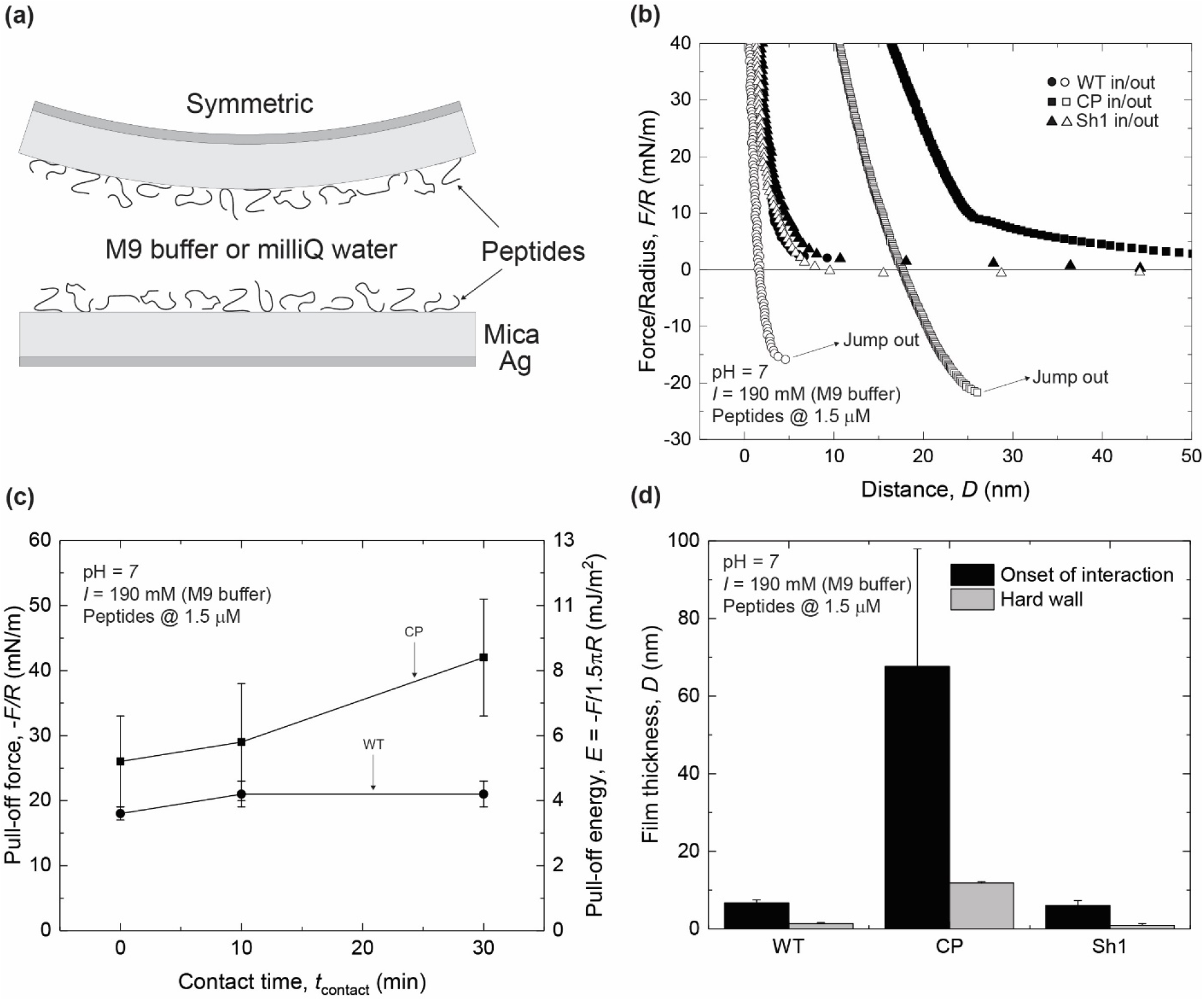
(a) SFA symmetric configuration used in these studies, consisting of back-silvered mica as substrates. Each mica supported a Bap1-57AA nanofilm deposited from a droplet of 40 µL at 1.5 µM, bridging the two surfaces. (b) Representative force-distance curves obtained during a loading and unloading cycle of WT (circles), CP (squares), and Sh1 (triangles) in M9 buffer. (c) Plots of pull-off forces normalized by radius of curvature, -*F/R*, versus contact time, *t*_contact_, for WT and CP peptides in M9 buffer. (d) Absolute film thicknesses at rest (onset of interaction) and at maximum compression (hard wall). Film thickness is reported as the thickness of the two films shown in the schematic (a).

It is well established that salts affect intra- and intermolecular forces, such as cation-π or hydrogen bonds.^34–36^ To investigate the adhesive and cohesive performance of the peptide films in the absence of salts, WT, Scr, CP, and Sh1 peptide films were deposited onto mica substrates from milliQ water, yielding a similar experimental configuration to that used for M9 buffer measurements (Figure 2a). Representative SFA force-distance curves in milliQ water, measured for WT, CP, and Sh1 are shown in Figure 3b. All force-distance curves for WT, Scr, and CP, exhibited purely repulsive forces during approach. Scr formed large aggregates, resulting in double contacts, and it was not possible to extract F-D profiles of the force-spectroscopy measurements performed with the SFA, similar to what observed for Scr in M9. Only CP had an abrupt jump out of contact during retraction of the surfaces. The pull-off forces, -*F/R*, for CP in milliQ water were 16.0 ± 1.6 mN/m, with no dwell time. Once again, leaving the surfaces under the maximum applied load (□100 mN/m) for an additional 10 min or 30 min affected CP film pull-off forces, increasing -*F/R* to 19 ± 3 mN/m and 22 ± 5 mN/m, respectively (Figure 3c), corresponding to an increase of □20% and □40% in pull-off forces magnitude. Similar to the M9 case, these observations suggest that intra- and intermolecular interactions responsible for the adhesive and cohesive properties of the CP nanofilm were established during the shorter contact times and increased with increasing contact times. Contrary to what measured in M9 buffer, WT and Sh1 peptide nanofilms in milliQ water were thicker than CP, and larger than those formed from M9 buffer, despite the negligible pull-off forces, suggesting that physical absorption is not the only factor determining the pull-off force.

**Figure 3.**
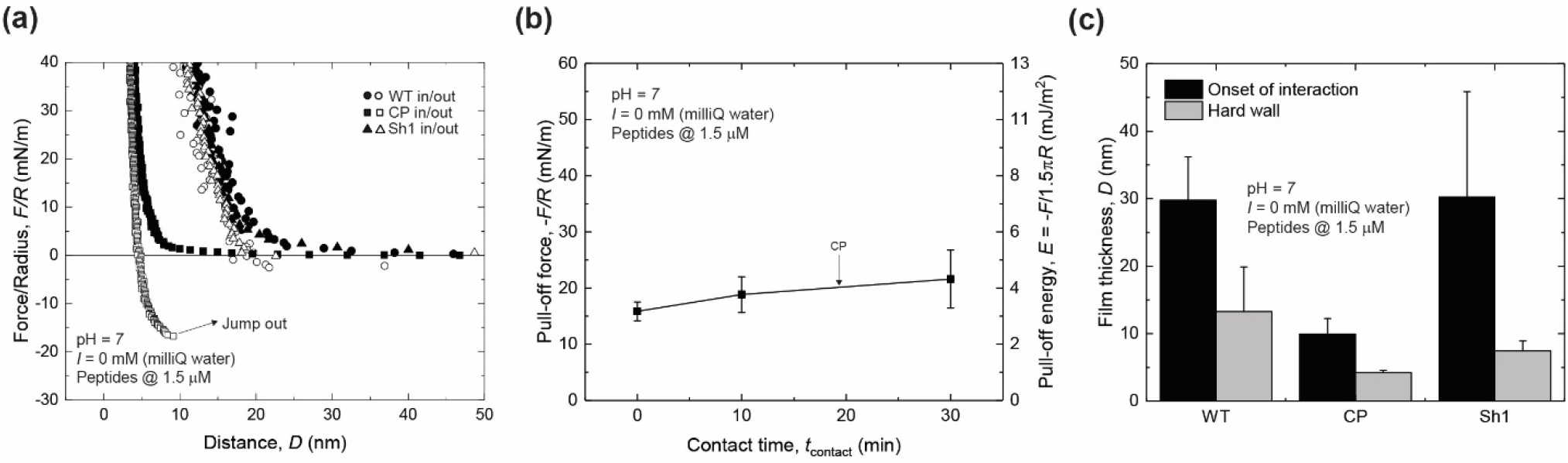
(a) SFA symmetric configuration representative force-distance curves obtained during a loading and unloading cycle of WT (circles), CP (squares), and Sh1 (triangles) in milliQ water. (b) Plots of pull-off forces normalized by radius of curvature, - *F/R*, vs contact time, *t*_contact_, for CP peptides in milliQ water. Film thickness is reported as the total thickness of compressed peptide films when the two surfaces approach, as shown in the schematic of Figure 2(a).

Lastly, we assessed the pull-off forces of WT, Scr, CP, and Sh1 peptides dissolved in DMSO, an organic solvent that is commonly used for solvating hydrophobic peptides, such as the ones investigated in this letter. A droplet at a bulk concentration of 1.5 µM of each peptide was deposited in a symmetric configuration, similar to the configuration used for M9 and milliQ water (Figure 2a). Purely repulsive forces were measured for WT, CP, and Sh1 during loading (approach), and no jump-outs of contact during retraction were detected (Figure S3). For the Scr peptide, no pull-off forces were collected due to the consistently large aggregates resulting in double contacts and long-range repulsion.

We now put our results into the context of existing literature on underwater adhesives, particularly those based on mfps. Mfp-3 and Mfp-5, two of the mfps with the highest catechol content, have 21% and 27% dihydroxyphenylalanine (DOPA).^37,38^ Higher contents of DOPA, as tested in synthetic constructs, such as PEGtides^34^ and peptides,^39^ do not increase the wet adhesion performance, indicative that other intra- and intermolecular interactions synergize for optimal surface bridging. The *V. cholerae* Bap1-inspired peptides presented here, specifically WT and CP, outperformed mfp-5 in very similar experimental conditions, with pH being the most significant difference.^40^ Particularly, CP peptide nanofilms formed from aqueous solutions with similar bulk molarity onto mica substrates in a high ionic strength environment at neutral pH, were more than 3× stronger for 2-minute contact times than mfp-5 nanofilms (25 mN/m for CP vs. 8 mN/m for mfp-5). We also note that the pull-off forces of CP nanofilms could potentially be further improved using longer dwelling times. Furthermore, the peptides used in this study consist exclusively of canonical amino acids (no DOPA). The performance of CP is partially attributed to its composition: 50% aromatic-rich amino acids, 40% content of polar amino acids, and 10% cationic content (Figure 1b). Based on back-of-the-envelope calculations, each CP peptide could form a maximum of 10 possible π-π bonds, two possible cation-π interactions, and five hydrogen bonds (donors or acceptors) with a surface or another peptide. Details on the estimates can be found in S8 Estimation of intermolecular interactions. The final, optimal number will be a function of the peptide’s conformational flexibility, side chain orientation, and environmental factors, such as the pH value and the nature of the ions present, as in the case of the M9 buffer, which is rich in phosphates. On the other hand, sequence also matters: the WT Bap1 57AA shows considerable pull-off forces, while the Scr version with the same composition has no measurable force. The contribution of the secondary structure of the peptide to adhesion will be an interesting topic for future study.

A challenge encountered was the frequent presence of large aggregates that made SFA measurements difficult. On average, one out of three independent experiments (one independent experiment consisted of a freshly prepared pair of mica substrates) was successful. To quantify aggregate sizes, dynamic light scattering (DLS) was used, and confirmed the presence of large aggregates (Figure S4 and S9. Peptide hydrodynamic size measured via dynamic light scattering), with estimated average hydrodynamic size for WT and CP aggregates larger than 1 µm when dissolved in M9 buffer, decreasing to hundreds of nm when dissolved in milliQ water. This is not surprising, as, in addition to the rich hydrophobic content, the presence of diphenylalanine (FF) in the sequence of the WT and CP was likely an additional driver for self-assembly, known to participate in β-amyloid-like fibril formation.^41^

In summary, the underwater cohesion and adhesion mechanisms of *V. cholerae* inspired peptides were studied using the SFA. For the first time, direct force spectroscopy of peptide sequences that mediate biofilm adhesion was investigated to identify new molecular motifs, based on canonical amino acids, that will aid in determining molecular design rules for robust underwater adhesives at neutral pH and in the presence of ions. The CP peptides exhibited excellent cohesive energy, reaching approximately 8 mJ/m^2^ in M9 buffer and approximately 4.5 mJ/m^2^ in milliQ water. Our results emphasize the importance of π-π stacking and cation-π, in addition to hydrogen bonding for the design of wet adhesives.

## AUTHOR INFORMATION

### Author contributions

The manuscript was written with contributions from all authors. All authors have approved the final version of the manuscript.

## ACKNOWLEDGEMENTS

S.T.A., S.Y.J. and X.H. acknowledge Z., A.I., A.D.M., and R.C.A.E. acknowledge funding from the National Science Foundation (NSF)-CREST: Center for Cellular and Biomolecular Machines through the support of the NSF Grant No. NSF-HRD-1547848. R.C.A.E. and S.T.A. acknowledge funding from the Defense Advanced Research Projects Agency through the GLURE: Grip Likelihood in Underwater Environments Grant No. HR0011-24-3-03-62 awarded to R.C.A.E.. funding from the Defense Advanced Research Projects Agency through the GLUE: Grip Likelihood in Underwater Environments Grant No. HR0011-24-3-03-62 awarded to J.Y. The views, opinions, and/or findings expressed are those of the authors and should not be interpreted as representing the official views or policies of the Department of Defense or the U.S. Government. J.Y. acknowledges support from the National Institutes of Health (NIH) Grant No. DP2GM146253. Additionally, S.Z. acknowledges funding from the NIH G-RISE I-BioSTeP Grant No. T32 GM141862. A.D.M acknowledges funding from the NSF Grant No. CHE-2316870.

## ABBREVIATIONS

SFA: surface forces apparatus
mfps: mussel foot proteins
WT: wild type
Scr: scrambled
Sh1: short 1
CP: central portion
DOPA: dihydroxyphenylalanine.

## Notes

### Competing Interest Statement

The authors have declared no competing interest.

