## Supplemental Information for "*Vibrio cholerae* adhesin-derived peptide mediates strong pull-off forces in aqueous high-ionic-strength environments"

**S1.** Bap1-inspired peptide details
**Table S1.** Library of Bap1-inspired peptides.
**S2.** Peptide synthesis and characterization.
**Fig. S1.** Mass spectra and corresponding HPLC traces
**S3.** Buffer receipt
**Table S2.** Adjusted M9 buffer salt concentrations.
**S4.** Surfaces Forces Apparatus (SFA) and substrate preparation
**S5.** Peptide film depositions
**S6.** Statistical analysis
**Table S3.** Summary of SFA data of CP in M9
**Table S4.** Summary of SFA data of CP in milliQ water
**Table S5.** Summary of SFA data of WT in M9
**Figure S2.** (a) Representative normal forces $F$ between negatively charged mica surfaces as a function of the mica-mica distance $D$ with a 50 μL droplet of M9 buffer
**S7.** Pull-off forces of Bap1-inspired peptides in DMSO
**Figure S3.** Representative normal forces normalized by radius of curvature $F/R$ between negatively charged mica surfaces with peptides films formed from a bulk concentration of 1.5 mM in DMSO
**S8.** Estimation of intermolecular interactions
**Figure S4.** Molecular structure of CP used to estimate π-π stacking, cation-π interactions, and hydrogen bonding, excluding backbone intermolecular interactions.
**S9.** Peptide hydrodynamic size measured via dynamic light scattering (DLS)
**Figure S5.** Hydrodynamic diameter bar plot of peptides in M9 and milliQ water
**Table S6.** Average and standard error of the mean data of hydrodynamic diameter of peptides in M9 and milliQ water

**S1. Bap1-inspired peptide details.** The 57-amino acid peptide found in Bap1 has a molecular weight of 6,793 Da. The wildtype Bap1 (WT) peptide amino acid (AA) sequence is (YLGLEWKTKTVPYLGVEWRTKTVSYWFFGWHTKQVAYLAPVWKEKTIPYAVPVTLSK). This sequence corresponds to the AA residues 415 to 471 from the Bap1 (or hemolysin) adhesin of *Vibrio cholerae*. The scrambled Bap1 (Scr) peptide AA sequence investigated in this study is (KWPGVSQYTLGKVVSLLTATLYYPEPEAKYVWKAWVTTLFYVGTERKKTIVWPFKWH). It was generated using the shuffle protein function from bioinformatics.org (accessed on 011/09/2024). Two additional shorter peptides, referred to in this study as short 1 (Sh1) and the central portion (CP), were synthesized as described below. The sh1 peptide sequence is WKTKTVPY, with a molecular weight of 1022 Da. The CP peptide sequence is SYWFFGWHTK, with a molecular weight of 1359 Da. Details of the peptide sequences are summarized in Table 1.

**Table S4.** Library of Bap1-inspired peptides.

| Peptide | Sequence | Molecular weight, MW (Da) | Solvents tested |
|---|---|---|---|
| WT | YLGLEWKTKTVPYLGVEWRTKTVSYWFFGWHTK QVAYLAPVWEKTIPYAVPVTLSK | 6793 | Water, M9, DMSO |
| Scr | KWPGVSQYTLGKVVSLLTATLYYPEPEAKYVWKA WVTTLFYVGTERKKTIVWPFKWH | 6793 | Water, M9, DMSO |
| Sh1 | WKTKTVPY | 1022 | Water, M9, DMSO |
| CP | SYWFFGWHTK | 1359 | Water, M9, DMSO |

**S2. Peptide synthesis and characterization.** Bap1-inspired peptides (WT, Scr, sh1, and CP) were either chemically synthesized and purchased from Atlantic Peptides (New Hampshire, USA) or LifeTein (New Jersey, USA). A second batch of each construct was synthesized in our laboratories, using the reagents and protocol listed below. Unless otherwise specified, all chemical reagents were sourced from Sigma Aldrich (St. Louis, MO) or ThermoFisher Scientific Inc. (Waltham, MA). Fmoc amino acids, Oxyma Pure, and resins were obtained from CEM. Matrix-Assisted Laser Desorption Ionization Time-of-Flight (MALDI-TOF) mass spectrometry data were acquired using a Bruker Microflex LRF mass spectrometer (positive reflector mode), with α-cyano-4-hydroxycinnamic acid (CHCA) as the ionization matrix (1:1 sample to matrix ratio). Purified peptides were lyophilized and later weighted using a Mettler Toledo XSR105DU analytical balance, to prepare stock solutions. Isolated peptides from reverse-phase HPLC were dialyzed against 1 liter of HPLC grade water in a 2,000 MWCO Slide-A-Lyzer Dialysis® cassette.

Peptides were synthesized using a CEM Liberty Blue microwave-assisted automated peptide synthesizer. Standard Fmoc Solid-Phase Peptide Synthesis (SPPS) involved iterative deprotection and coupling cycles. Deprotection was carried out with 4-methylpiperidine (4MP) in dimethylformamide (DMF; 20% v/v), while coupling was mediated by DIC/Oxyma Pure activation. Peptide synthesis was performed on Cl-TCP(Cl) ProTide resin (CEM), with both the C- and N-termini left unmodified. Double-coupling cycles were used after coupling 30 amino acids. Crude peptidyl resins were filtered, rinsed with acetone, and air-dried. The peptides were then cleaved from the resin for 3 hours at room temperature using a cleavage mixture consisting of 92.5% trifluoroacetic acid (TFA), 2.5% triisopropylsilane (TIPS), 2.5% 3,6-dioxa-1,8-octanedithiol (DODT), and 2.5% $H_2O$. The cleaved peptides were precipitated with cold diethyl ether (5°C) and centrifuged at 4,500 rpm for 15 minutes at 5°C. The supernatants were discarded, and the resulting pellets were dried under vacuum overnight. Due to poor solubility, crude peptides were prepared in acetonitrile and water (45:55, respectively) with 0.1% TFA. Peptides were purified by reverse-phase HPLC using an Agilent 1260 Infinity II prep-scale HPLC system, equipped with a Kinetex XB-C18 column (250 × 30 mm, 5 µm, 100 Å). An additional HPLC purification run was carried out for the wild-type bacterial adhesin peptide (WT Bap1).

The peptides were eluted using a linear gradient of acetonitrile and water containing 0.1% TFA. Target fractions containing the desired peptide (confirmed by MALDI-TOF mass spectrometry) were collected, concentrated by rotary evaporation, lyophilized. The lyophilized peptides were dialyzed against $H_2O$) to remove residual TFA (MWCO = 2000 Da). The resulting peptide solutions were lyophilized once more and stored at -30°C. Purified peptides were analyzed via analytical HPLC analysis using an Agilent Infinity II 1260 analytical HPLC system, equipped with an Agilent Poroshell 120 EC-C18 column (100 × 2.7 mm, 2.7 µm, 120 Å). The HPLC run started with 30% acetonitrile in water and was held for 1 minute, after which the acetonitrile concentration was increased by 1.2% per minute until it reached 60%. Peak identification was achieved by collecting fractions using an automatic fraction collector and confirming peptide identity with MALDI-TOF mass spectrometry (Fig. S1).

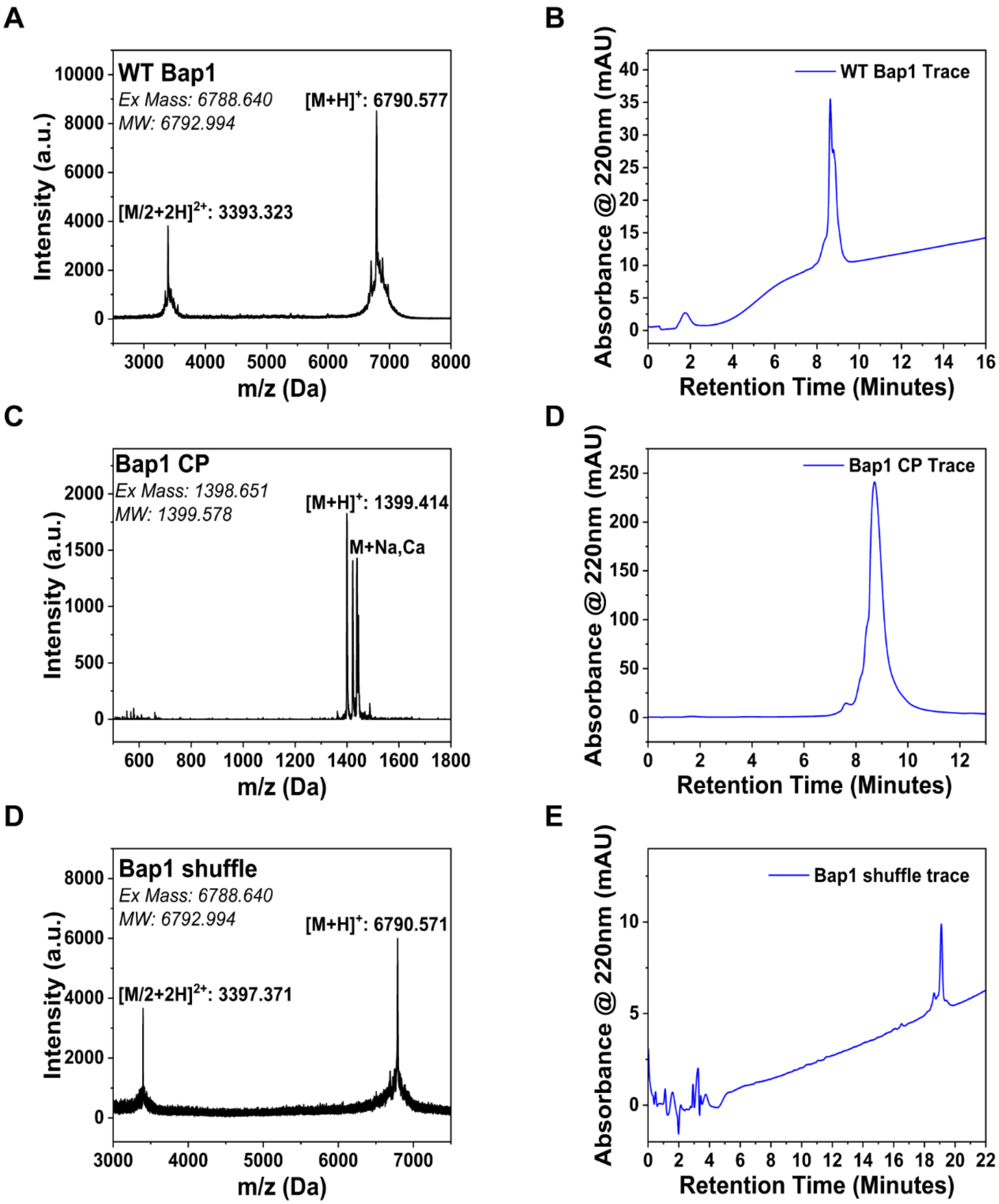


**Fig. S1.** Mass spectra and corresponding HPLC traces of WT (A) and (B), CP (C) and (D), and Scr (E) and (F).

**S3. Buffer receipt.** M9 buffer was made from M9 minimal salts, 5X (Sigma, Product # M6030), 1 M magnesium sulfate, $MgSO_4$ (J.T. Baker, Product #2500-01), 1 M calcium chloride, $CaCl_2$ (J.T. Baker, Product #1313-01) in ultrapure water (Millipore Sigma, resistance (18.2 MΩ-cm), resulting in a buffer of pH 7 and ionic strength of 192 mM, Table S2. The lyophilized peptide was dissolved to a stock concentration of 150 µM in dimethyl sulfoxide (DMSO, ≥ 99.9 % purity, Sigma) and stored at -20 °C until use. For dynamic light scattering (DLS) and Surface Forces Apparatus (SFA) measurements, the stock solution was diluted with M9 buffer or PBS buffer prefiltered using 0.2 µm filters (Millipore Sigma, MillexTM hydrophilic PTFE syringe filter, Product # SLLGC13NL), or milliQ water to a desired final concentration of 0.75, 1.5, and 8.5 µM and stored at 4 °C until use. All peptide solutions contained less than 6 vol. % DMSO.

**Table S5.** Adjusted M9 buffer salt concentrations.

| Salt Type | $KH_2PO_4$ | NaCl | $Na_2HPO_4$ | $NH_4Cl$ | $MgSO_4$ | $CaCl_2$ |
|---|---|---|---|---|---|---|
| [mM] | 22.1 | 8.6 | 48.0 | 18.7 | 2.0 | 0.1 |

**S4. Surfaces Forces Apparatus (SFA) and substrate preparation.** High-purity silver shots were purchased from Ted Pella (Product # 25-50, California, USA) and used to thermally evaporate ~ 50 nm of silver using a benchtop thermal evaporator (Denton Vacuum, NJ, USA) on freshly cleaved and hot Pt-wire precut mica (S&J Trading Inc., New York, USA) of thicknesses 3-4 µm. The back silvered mica surfaces were then glued with an epoxy resin (EPON 1004F, Miller-Stephenson) onto semi-cylindrical fused silica disks with radius of curvature, $R$ ~ 2 cm (Esco optics, USA), and arranged in the Surface Forces Apparatus (SFA 2000, SurForce LLC) in a crossed-cylinder geometry to construct the optical interferometer, which, combined with white light, gives rise to fringes of equal chromatic order (FECO). Details are described elswhere.[1,2] The lower surface was suspended on a double cantilever spring with a spring constant of $k_{spring}$ ~1,000 N/m. FECO were used to measure the absolute separation distances $D$ between the two mica surfaces. $D$ = 0 nm or reference distance was established from the contact of mica-mica in air (*i.e.*, without peptide films) for every new set of mica substrates. Normal force $F$ was obtained by subtracting the measured position of the surfaces from the expected position of the surfaces in the absence of a force field, corresponding to the spring deflection. Force $F$ was normalized by the radius of curvature $R$. Positive $F/R$ values correspond to repulsive forces. Negative $F/R$ corresponds to attractive (adhesive and cohesive) forces. The radius of curvature is the equivalent radius of curvature $R$ resulting from the two semi-cylindrical surfaces, $1/R = 1/R_1+1/R_2$, mapped with FECO. FECO positions as a function of time were recorded with a Hamamatsu ORCA Flash scientific camera at a rate of 0.5 fps. Recorded timeframes were analyzed with a custom-written MATLAB script, and distances (and corresponding forces) were calculated using analytical solutions for a two-layer optical interferometer.[3]

**S5. Peptide film depositions.** A symmetric, two-peptide film configuration was used in this study. The configuration consists of a symmetric peptide deposition achieved as follows: A 50 µL droplet containing the working concentration of Bap1-peptides of either DMSO, milliQ water, or M9 buffer was injected between two mica surfaces. The peptides were allowed to physisorb for 30 min at room temperature inside the SFA box, sealed, ending with a rinsing step using the incubation solvent to remove unbound peptides, leaving 30 μL between the surfaces for measurements. During each measurement, the surfaces were brought into contact at ~15 nm/s, compressed to $F/R$~100 mN/m, and held in contact for 2, 10, or 30 min, and then separated at a velocity of ~ 15 nm/s using a motorized differential micrometer with a reduction of 1670:1. The maximum attractive force measured during separation was denoted as the pull-off force, $-F/R$. Pull-off energies $E$ were calculated using the Johnson–Kendall–Roberts (JKR) theory: $E = -F/1.5\pi R$.[4] During force measurements, a glass boat with ultrapure water was placed inside the SFA chamber to maintain a saturated vapor environment and minimize meniscus evaporation between the surfaces. Tables S3, S4, and S5 provide averages and standard error of the means (SEM) for the maximum pull-off forces, $-F/R$, jump-out distance, $D_{\text{jump-out}}$, onset of interaction, $D_0$, and hardwall, $D_{\text{HW}}$. For the conditions in which pull-off forces were measured. That is, CP in M9, CP in milliQ water, and WT in M9, respectively.

**S6. Statistical analysis.** SFA force spectroscopy results are reported as the averages and standard error of the means from experiments performed with at least two independent sets of surface pairs, with three force-distance measurements per position and two contacts per set of surface pairs. DLS results are reported as the averages and standard error of the mean performed with at least two independent samples, with three measurements per sample

**Table S6.** Summary table of averages and standard error of the means (SEM) for the maximum pull-off forces, $-F/R$, jump-out distance, $D_{\text{jump-out}}$, onset of interaction, $D_0$, and hardwall, $D_{\text{HW}}$ collected from SFA measurements for CP in M9.

| **Contact time, $t_{\text{contact}}$ (min)** | **Maximum pull-off forces, -*F/R* (mN/m)** | **Jump-out distance, $D_{\text{jump-out}}$ (nm)** | **Onset of interaction, $D_0$ (nm)** | **Hardwall, $D_{\text{hardwall}}$ (nm)** |
|---|---|---|---|---|
| **0** | -26 ± 7 | 30.0 ± 3.0 | 70.0 ± 30.0 | 11.8 ± 0.5 |
| **10** | -29 ± 9 | 36.0 ± 9.0 | 70.0 ± 20.0 | 12.7 ± 0.5 |
| **30** | -42 ± 9 | 50.0 ± 20.0 | 80.0 ± 20.0 | 20.0 ± 9.0 |

**Table S4.** Summary table of averages and standard error of the means (SEM) for the maximum pull-off forces, $-F/R$, jump-out distance, $D_{\text{jump-out}}$, onset of interaction, $D_0$, and hardwall, $D_{\text{HW}}$ collected from SFA measurements for CP in milli-Q water.

| **Contact time, $t_{\text{contact}}$ (min)** | **Maximum pull-off forces, -*F/R* (mN/m)** | **Jump-out distance, $D_{\text{jump-out}}$ (nm)** | **Onset of interaction, $D_0$ (nm)** | **Hardwall, $D_{\text{hardwall}}$ (nm)** |
|---|---|---|---|---|
| **0** | -10 ± 3 | 9.5 ± 2.0 | 10.5 ± 2 | 4.3 ± 0.5 |
| **10** | -10 ± 4 | 7.0 ± 1.5 | 10.6 ± 2 | 4.5 ± 0.5 |
| **30** | -10 ± 5 | 8.0 ± 2.0 | 10.5 ± 2 | 4.3 ± 0.5 |

**Table S5.** Summary table of averages and standard error of the means (SEM) for the maximum pull-off forces, -*F*/*R*, jump-out distance, $D_{jump\text{-}out}$, onset of interaction, $D_0$, and hardwall, $D_{HW}$ collected from SFA measurements for WT in M9.

| **Contact time, $t_{contact}$ (min)** | **Maximum pull-off forces, -*F/R* (mN/m)** | **Jump-out distance, $D_{jump\text{-}out}$ (nm)** | **Onset of interaction, $D_0$ (nm)** | **Hardwall, $D_{hardwall}$ (nm)** |
|---|---|---|---|---|
| **0** | -18 ± 1 | 5.2 ± 0.5 | 6.4 ± 0.5 | 1.1 ± 0.5 |
| **10** | -21 ± 1 | 6.0 ± 2.0 | 6.0 ± 2.0 | 0.8 ± 0.5 |
| **30** | -21 ± 2 | 4.5 ± 1.0 | 4.6 ± 0.5 | 0.8 ± 0.5 |

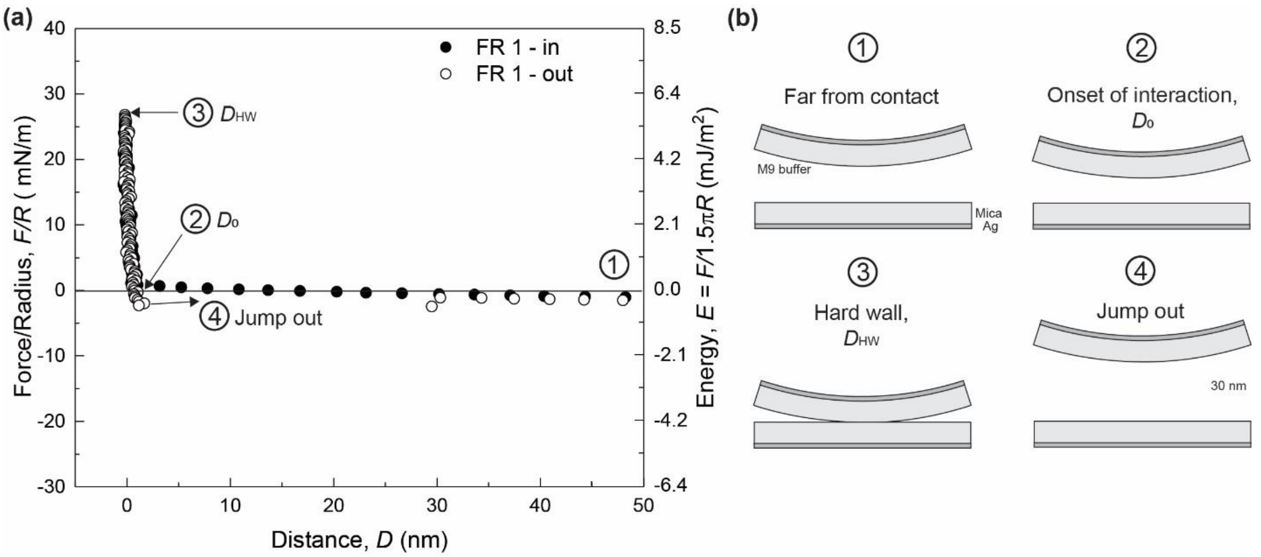


**Figure S2.** (a) Representative normal forces *F* between negatively charged mica surfaces as a function of the mica-mica distance *D* with a 50 µL droplet of M9 buffer. The force *F* is normalized by the radius of curvature *R* of the mica surfaces. Solid and open symbols indicate approaches and retractions, respectively. (b) Schematics indicate the initial surfaces at a finite distance (1), the onset of interaction, or distance at which the two nanofilms begin to interact (2), hard wall or film thickness at maximum compressive force (3), and separated surfaces after jumping out of contact (4).

### S7. Pull-off forces of Bap1-peptdies in DMSO

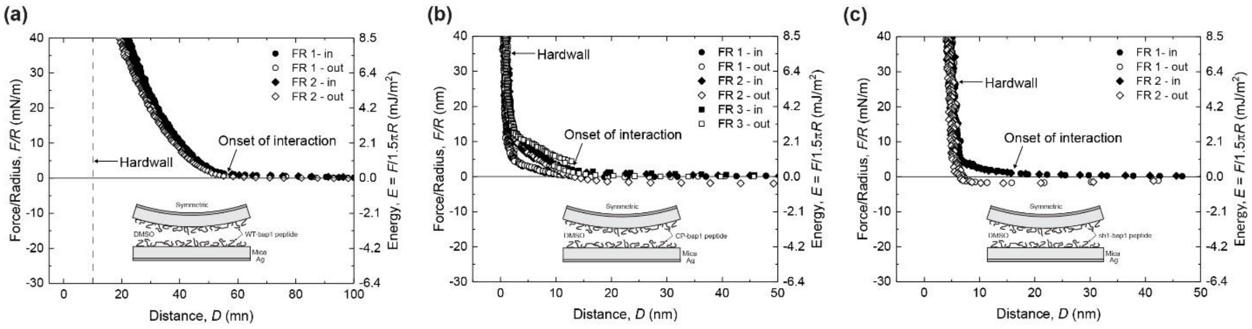


**Figure S3.** Representative normal forces normalized by radius of curvature *F/R* between negatively charged mica surfaces with peptides films formed from a bulk concentration of 1.5 µM in DMSO for (a) WT, (b) CP, and (c) Sh1 as a function of the mica-mica distance *D* with a 40 µL droplet of DMSO. Solid and open symbols indicate approaches and retractions, respectively.

### S8. Estimation of intermolecular interactions in CP peptide

#### π-π stacking

Residues that have the potential to participate in π-π stacking from the CP peptide include tyrosine (Y), tryptophan (W), phenylalanine (F), and histidine (H), which form weaker π-π interactions. Tryptophan and phenylalanine appear twice in the sequence, and excluding histidine, the sequence ends with a total of five aromatic residues. With two interactions possible per residue, there are a maximum of 10 unique pairwise interactions.

#### Cation-π interactions

In addition to the five residues listed above, histidine (H) and lysine (K) contribute as cations in the sequence. Taking the minimum value between the π-system donors (five) and the two cations, the total maximum of π-cation interactions that CP can form is estimated to be two.

#### Hydrogen bonding

From the sequence of CP, serine, tyrosine, tryptophan, histidine, and threonine can participate as hydrogen bond acceptors. Lysine, depending on its protonation state, can participate in one additional hydrogen bond, yielding 5-6 hydrogen bonds.

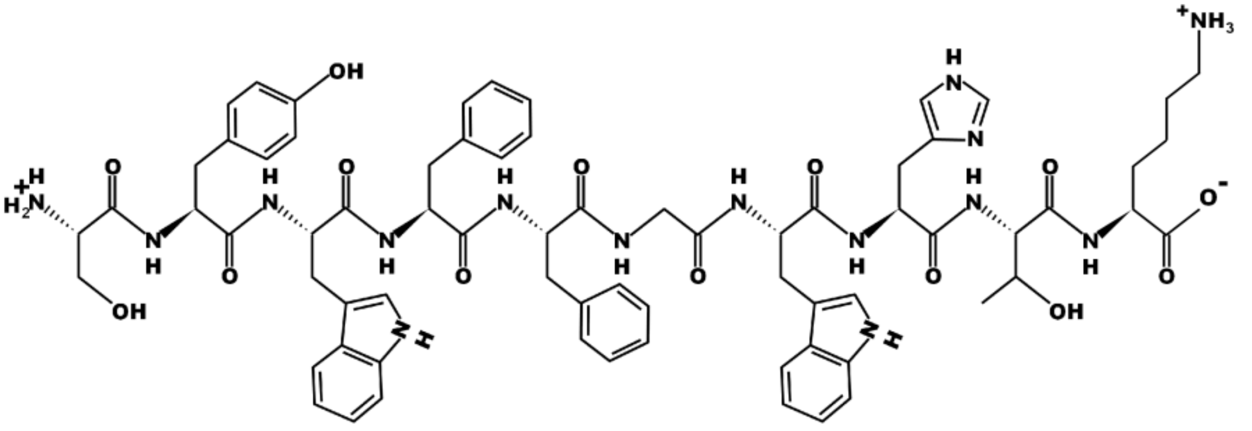

**Figure S4.** Molecular structure of CP used to estimate Pi-Pi stacking, Pi-cation interactions, and hydrogen bonding, excluding backbone intermolecular interactions.

### S9. Peptide hydrodynamic size measured via dynamic light scattering (DLS)

All solvents were filtered through a 0.2 μm hydrophilic PTFE syringe filter before use. Stock solutions of 1200 μM peptides (WT and Scr) were prepared in DMSO and acetonitrile by dissolving the peptide powders in the filtered solvent within 1.5 mL low-binding microcentrifuge tubes. The peptide solubility was assessed by observing potential aggregation after pipetting the solution up and down several times (Table I). In DMSO, both peptides were fully dissolved and formed clear solutions. However, attempts to dissolve peptides in acetonitrile under the same conditions produced cloudy suspensions.

To further study peptide solubility in aqueous-based solvent mixtures, peptide-DMSO stock solutions were diluted to 1.5 μM using either adjusted M9 buffer (with high ionic strength of 190 mM), 1X PBS (pH 7.4), and ultrapure water (ThermoFisher). A series of dilutions (75 μM, 37.5 μM, 8.5 μM, and 1.5 μM) in M9 were also performed. At higher concentrations (75 μM and 37.5 μM), both peptides formed cotton-like aggregates, while no aggregation was observed at 8.5 μM and 1.5 μM. At 1.5 μM, all peptide solutions were visually clear, likely due to low concentration.

To examine solubility at 1.5 μM in greater detail, hydrodynamic particle size was measured using Dynamic Light Scattering (DLS). DMSO-M9, DMSO-PBS, and DMSO-$H_2O$ solutions showed increased particle size in salt-containing conditions, suggesting salts negatively affected peptide dispersion or promoted aggregation.

In a separate test, the 1200 μM peptide-acetonitrile solutions were stored at 4°C for three weeks to observe potential changes over time. No dissolution was observed; instead, most of the acetonitrile evaporated. After storage, adding fresh acetonitrile to the WT sample to achieve a 200 mM concentration resulted in the formation of visible flakes. It is possible that undissolved peptides crystallized during solvent evaporation over the prolonged storage period. To test whether the crystals could be redissolved, DMSO was added to the Scr sample post-evaporation. The resulting clear solution indicated that DMSO is a highly effective solvent for both 57AA peptides, outperforming acetonitrile, especially in redissolving peptides in crystallized forms.

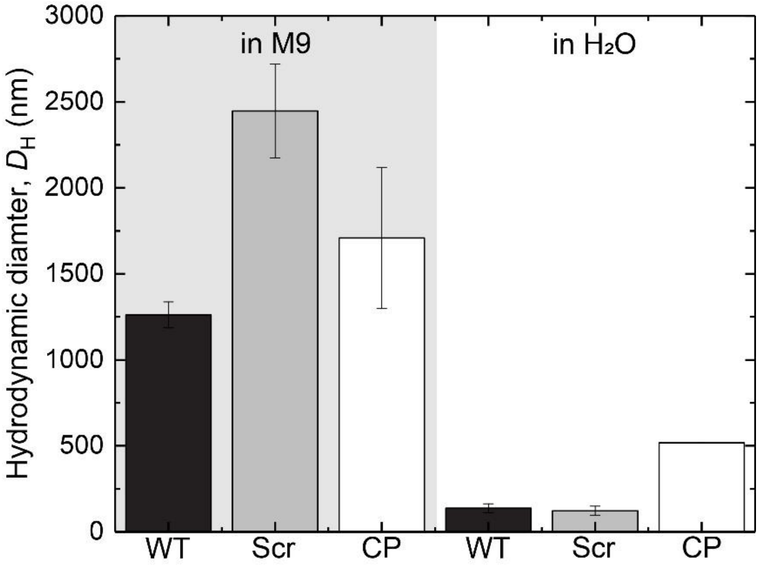


**Figure S5.** Hydrodynamic diameter of (a) WT-Bap1 and (b) Scr-Bap1 peptide in DMSO at 150 μM. In water, PBS, and M9, the concentration of (a) WT-Bap1 and (b) Scr-Bap1 peptide is 1.5 μM, used for the deposition in SFA pull-off force measurements, corresponding to what is referred in the main text as intermediate bulk concentration. Note differences in the hydrodynamic diameter scale between WT-Bap1 and Scr-Bap1. The average ± standard error of the mean is shown from at least two independent measurements.

**Table S6.** Summary table of averages and standard error of the means (SEM) for the maximum pull-off forces, $-F/R$, jump-out distance, $D_{jump\text{-}out}$, onset of interaction, $D_0$, and hardwall, $D_{HW}$ collected from SFA measurements for CP in M9.

| Peptide | Average hydrodynamic diameter in M9, $D_H$ (nm) | Average hydrodynamic diameter in $H_2O$, $D_H$ (nm) |
|---|---|---|
| **WT** | 1261.5 ± 75.5 | 137.3 ± 25.5 |
| **Scr** | 2446.5 ± 273.5 | 122.9 ± 26.0 |
| **CP** | 1709.0 ± 410.0 | 518.0 ± 1.3 |
